# The Untangle Challenge for accurate ensemble models

**DOI:** 10.64898/2026.02.21.706873

**Authors:** Mehagan S. Hopkins, Thomas C. Terwilliger, Pavel V. Afonine, Helen M. Ginn, James M. Holton

## Abstract

We report the discovery of a new class of local minima that has severely limited the accuracy of macromolecular models. Termed density misfit barrier traps, these minima explain much of the poor fit between macromolecular models and experimental data relative to that of smaller molecules: not just high R factors, but distorted chemical geometry. We postulated that proteins exist as an ensemble of conformations that each have good geometry, but refinement algorithms have been unable to converge to them due to a tangling phenomenon arising from these traps. To demonstrate, a synthetic ground truth data set was generated, consisting of a 2-member ensemble with excellent geometry. A series of starting models, each trapped in increasingly difficult local minima, were prepared, a unified validation score defined, and an open Challenge issued. This Challenge inspired algorithms for escaping such traps, and new programs have been released that are expected to substantially improve the accuracy of macromolecular ensemble models.

**Synopsis:** A synthetic 2-member conformational ensemble of a small protein and corresponding electron density data was generated to demonstrate how topological local minima hinder simultaneous agreement with density data and chemical geometry restraints in conventional structure refinement.

## 1. Introduction

Despite decades of hard-won improvements in refinement and modeling software, macromolecular models remain a poor fit to accurately observed X-ray data, with disagreement usually four times larger than the errors in the experimental data (Vitkup *et al*., 2002, Holton *et al*., 2014). Due to Parseval’s Theorem, this poor fit leads to proportionately large errors in density maps (Lang *et al*., 2010). It also distorts the chemical geometry of the model away from canonical values (Engh & Huber, 2012) seen in smaller structures. Both of these factors limit the interpretability of macromolecular models, and reconciling this tug-o-war (Urzhumtseva *et al*., 2009) between density and geometry will unlock hidden potential in the very high quality data that can be obtained from macromolecular crystals. Figure 1 shows how increased model accuracy lowers the noise floor of electron density maps. This reduction of phase errors has the potential to reveal low-lying features that are currently unobservable. Not just hydrogens, but other subtle changes, such as weakly-bound ligands (Liebschner *et al*., 2017, Pearce *et al*., 2017), low-occupancy-high-energy states (Lang *et al*., 2014, Burnley *et al*., 2012), and other gating conformations could become routinely visible, even at moderate resolution. This is because modern crystallographic data are accurate enough to achieve signal-to-noise ratios of 100 or higher (1% error) (El Omari *et al*., 2024), as is often required for anomalous phasing (Hendrickson & Teeter, 1981, Liu & Hendrickson, 2017, Terwilliger *et al*., 2016), but map accuracy is currently limited by Parseval’s Theorem to no more than signal-to-noise ∼5 because R factors are seldom better than 20%.

**Figure 1:**
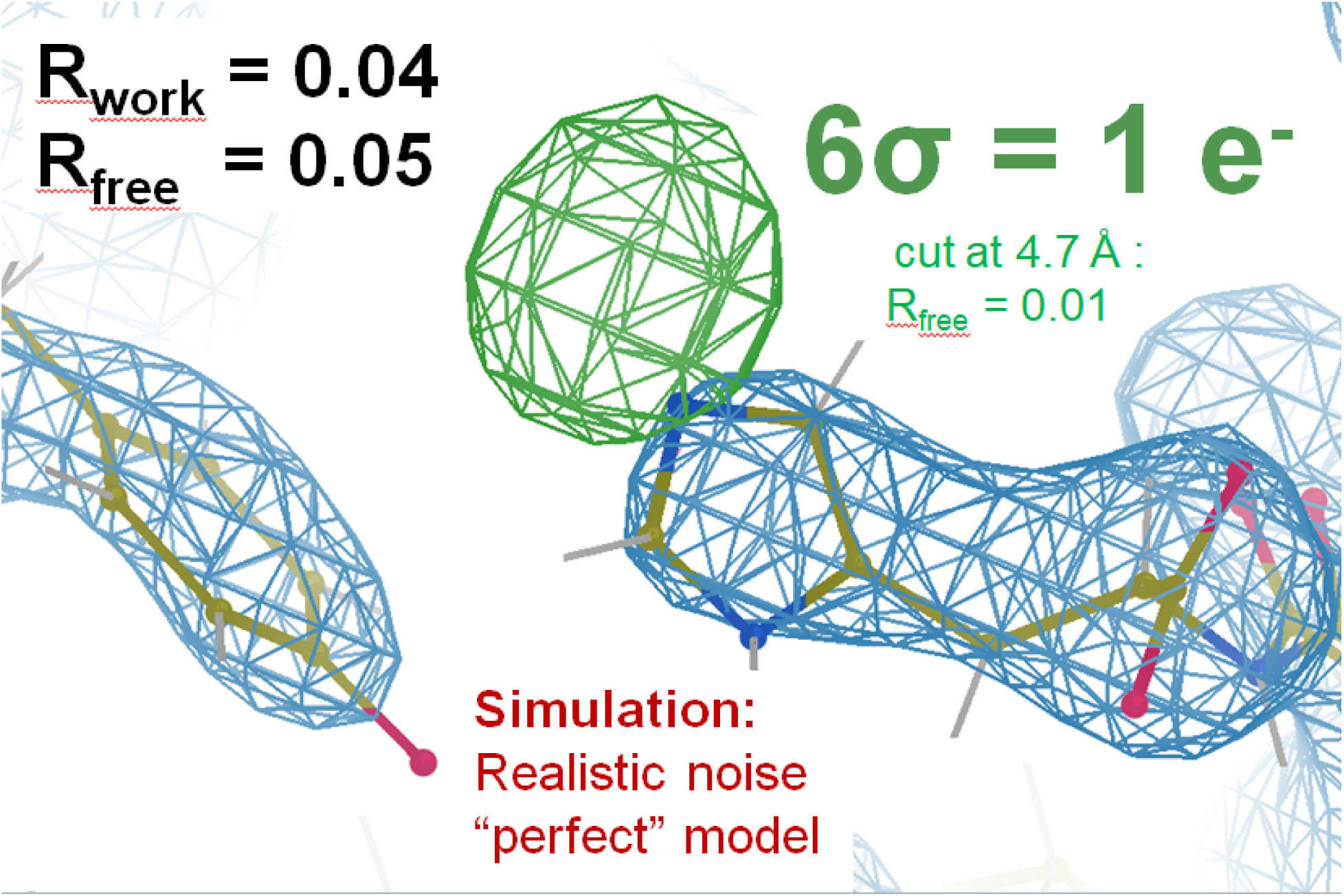
The potential power of low R-factors in revealing hidden features: One missing hydrogen atom generated a 9-σ peak, contoured at 6σ (green) using data cut to 4.5 Å resolution in an mF_sim_-DF_c_ map. We write F_sim_ instead of “F_obs_” because F_sim_ was obtained by processing diffraction images simulated with realistic experimental errors as described in Appendix A. Resolution limit where CC_1/2_=0.3 was 3.07 Å, but maps were cut further and sharpened to optimize H peak height. 2mF_sim_-DF_c_ map was contoured at 1σ and cut at 3.5 Å (blue). F_c_ was the ground truth of the image simulation after H removal and a single round of refmac5. Refinement of a single-copy model like the one shown for clarity yielded R_work_/R_free_ = 0.25/0.3 and no informative difference peaks (not shown). Although merely a simulation, this example demonstrates the remarkable sensitivity that is possible with X-ray diffraction data, provided model bias and all other systematic errors are removed. Experimental errors do not hide single-electron features, modelling errors do.

A popular explanation for high R factors and poor geometry is that protein structures, even at cryogenic temperatures, exist as ensembles, and these ensembles are imperfectly captured by the standard, largely single-conformer models used to represent them (Terwilliger *et al*., 2007). Over the decades, this hypothesis has been tested (Burling & Brünger, 1994, Burling *et al*., 1996, Vitkup *et al*., 2002, Wankowicz *et al*., 2024, Burnley *et al*., 2012) but the model-data agreement has never approached that of chemical crystallography. The low observation/parameter ratio of high-copy ensemble models has often been given as a reason for returning to the more parsimonious traditional representation.

Another common hypothesis is that high R factors are due to poor resolution of the data, but the simulation shown in Figure 1 demonstrates that experimental errors expected for this modest resolution data set do not have this effect on R_work_ and R_free_. Indeed, poor resolution lowers the observations/parameters ratio, ostensibly making the data easier to fit to a complex model, not harder. For example, if only a few hundred observations are available and the model has millions of parameters, one would expect R_work_ values near zero could be readily obtained, which is not the case in practice. This is because refinement programs use an energy function (Headd *et al*., 2012, Afonine *et al*., 2012) that penalizes distortions away from chemically reasonable bond lengths, angles, and other prior knowledge of inter-atomic geometry. Simultaneously satisfying this prior knowledge at the same time as the accurate X-ray data remains an elusive, yet fundamentally expected, outcome.

### 1.1 Energy models of chemical geometry

The refinement geometry energy is related to the real-world energy landscape of the molecule (Moriarty *et al*., 2020), with important differences: there are no electrostatics, non-bond interactions are softened and repulsive-only, and quantum-mechanical effects are neglected (Zubatyuk *et al*., 2025, Janowski *et al*., 2016). The geometry energy is usually a statistical potential (Moriarty *et al*., 2014), where expected bond lengths, angles and other geometric values are taken from the average of a database of high-quality structures, and the standard deviation of the observed distribution defines the width of a harmonic potential well about that mean.

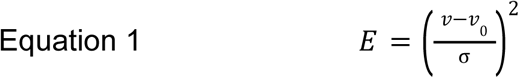

Where: E is the geometry energy, v_0_ is the ideal value of a bond length or other parameter, v is the value in the model, and σ is the root-mean-square from the database, taken as a standard deviation. It is noteworthy here that these standard deviations of bonds and angles are typically ∼0.02 Å, and ∼2° but can be as low as 0.005 Å and 1°. No one has ever collected a dataset to 0.005 Å resolution, and yet bond lengths are known to this level of accuracy. This makes geometry energy a powerful restraint, and the present work tested just how powerful.

An important difference between a statistical energy and chemical energy is that chemical energy reflects the number of Joules stored in the molecule by distorting it away from its ground state configuration, but a statistical energy reflects how often a given distortion is observed. This relationship is often compared to a Boltzmann distribution, but this analogy has caveats. The strain observed in crystal structures is not thermal in origin. Instead, it comes from context. The surrounding atoms in the crystal push a given feature, very reproducibly, into what may be an uncommonly-observed distortion. In recent years, it has become evident that improved context-based classification is needed for statistical potentials to better reflect true chemical energy, such as the conformation-dependent library (CDL) in phenix (Tronrud *et al*., 2010, Tronrud & Karplus, 2011, Moriarty *et al*., 2016), and why quantum refinement is expected to improve geometry even further (Zubatyuk *et al*., 2025, Liebschner *et al*., 2023).

But, in the present work, for simplicity, portability and accessibility reasons, the ground truth of this Challenge was made to agree well with the more traditional and broadly-available statistical energy libraries (Engh & Huber, 2012) implemented in popular refinement programs. We further hypothesized a situation where v_0_ was the same in the geometry energy and the chemical energy, but the σ values in the geometry energy were larger than the actual chemical energy wells, due to averaging over varied contexts. We therefore expect the geometry energy of individual ensemble members to be superior to that of typical structures.

Finally, and most relevant here, all refinement programs must balance the geometry energy against a pseudo-energy reflecting how well the model agrees with the observed density data. In practice, agreement with both energies is poorer than expected from the standard deviations, begging the question: with such high quality data on both sides, why is there always a tug-of-war between geometry and density?

### 1.2 Local Minima

The complexity of this minimization invites the obvious possibility that protein structural models are trapped in local minima. Many types of local minima that pit the geometry term against the density term are well-known, such as peptide bond flips (Croll, 2015, Williams *et al*., 2018), sequence registration errors (Terwilliger, 2003), and the rotameric states of Asn, Gln and His side chains (NQH flips) (Lunin *et al*., 2002, Word *et al*., 1999, Headd *et al*., 2009). All of these examples are errors that cannot be corrected by simple minimization, and automated methods for identifying and correcting them are now mature (Headd *et al*., 2009).

Here we report a new class of local minima that only occur in ensemble models refined against density data. The distances moved to escape these minima are small, but the impact on the geometry energy can be dramatic. Escaping them not only results in more chemically reasonable models, but allows the density fit to improve, eventually revealing previously hidden details, like in Figure 1.

In current practice, there are three main ways to represent the ensemble nature of proteins. The most common and most parsimonious, is the “multi-conformer model” where most atoms are modeled with a single XYZ location, and only when density is encountered that shows clear evidence of an ensemble, relevant atoms in the model are split into “alternate locations” (altlocs). Programs such as qFit (Wankowicz *et al*., 2024) and Coot (Emsley *et al*., 2010) produce such models by default. Another practice, most common in NMR structures, is to denote each ensemble member as a complete copy of the molecule in its own right, often giving each member of the ensemble the same occupancy, as is the default in phenix.ensemble_refine (Burnley *et al*., 2012) and Vagabond (Ginn, 2023). A third option is to explicitly symmetry-expand the ensemble chains so that each member occupies different asymmetric units of the unit cell (Mikhailovskii *et al*., 2024), or even a supercell (as in Appendix A). Unlike the other two representations, the members of these “supercell ensemble” models interact with each other via non-bonded contacts at crystal packing interfaces. All atoms are at full occupancy, allowing bulk solvent to differ for each ensemble member. It is very important to note here, however, that despite this variety of available representations, all ensemble models are equally vulnerable to the density-misfit barrier traps reported here. This is because density obeys crystallographic symmetry. For this reason, we will refer to all of the above representations as “ensemble models” here.

In this work we focus on a 2-member ensemble: A and B. In some instances, e.g. a disulfide bond with two well-defined conformations in the x-ray density, it is clear which version of each atom belongs in the same conformation, however this is the exception and not the rule. For atoms that are mis-assigned, current software is better at finding nearby local minima for the defined conformer assignments than it is at reversing the swap, allowing models of incorrect correlated motions to survive refinement.

Here we refer to the new type of local minima that hinders all types of ensemble refinement as a density misfit barrier trap. Much in the same way that wires in a multi-conductor cable cannot be easily re-arranged inside a tight cable jacketing, the electron density cloud also bundles the chains of an ensemble model together. Once built, ensemble members cannot easily swap places because when they pass through each other the density fit is poor (Figure 2). Moving ensemble members to different asymmetric units, such as in supercell ensemble refinement (Mikhailovskii *et al*., 2024), does not remove these barriers because the density obeys crystallographic symmetry.

**Figure 2:**
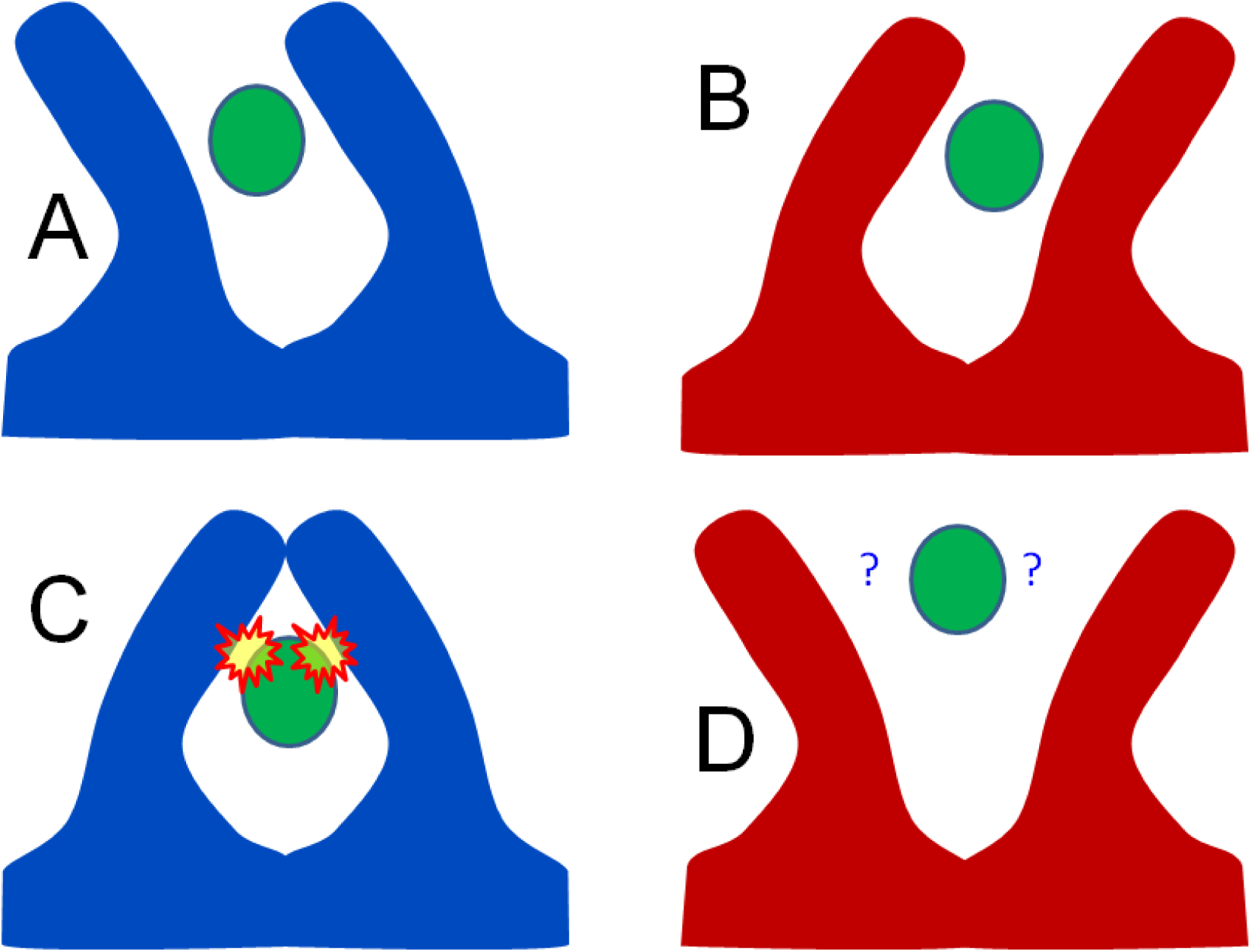
Schematic of how correlated vs anti-correlated motion can impact ligand binding. If the macromolecule conformers (red and blue) comprise a “windshield wiper” or correlated mode (A vs B), a ligand (green) bound between the two waving appendages will have similar contacts in both A and B conformations, and may bind tightly without clashes. On the other hand, if the molecular motion is anti-correlated, like “jumping jacks” (C vs D), then C will have bad clashes with the ligand and D will have loose contacts, limiting interactions. In this way, the mode of preferred macromolecular motion can have a dramatic impact on affinity. However, the average electron density of both A+B and C+D parings are identical. A single-conformer model built into that average density, even with anisotropic B factors, will be a poor proxy for the underlying situation. A 2-conformer model, however, can capture the molecular motion, but only if the conformer assignments are correct.

**Figure 3:**
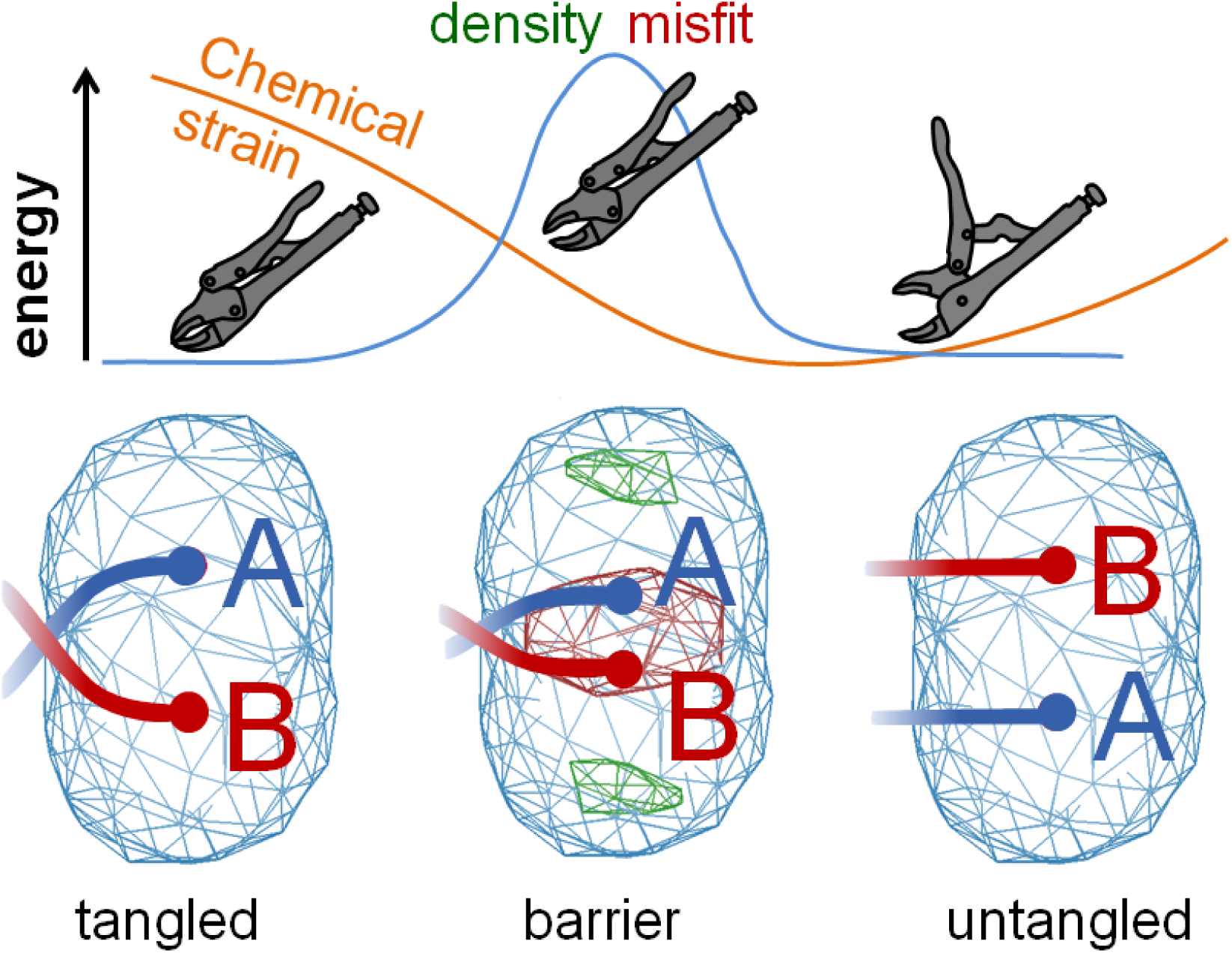
Simplest possible example of a density misfit barrier trap: The force that holds ensemble structures in high-energy local minima like this arises from the density term (blue line). The geometry term is represented by the orange line. The connectivity with the rest of the structure makes the B/A conformer assignment on the right more energetically favorable than the A/B assignment on the left. Despite this clear advantage on the right, the favorable state is unreachable because of the poor density fit at the midpoint of the transition. This is because as the two alternate atom locations pass through one another they form a high-density shape that is a poor match for the observed, oblong density cloud. Note the red and green difference density in the middle barrier state and not in the tangled and untangled states. The clamped, over-center, and released positions of locking pliers are shown to illustrate a mechanical analogy to such traps. In this way, density disagreement forms a barrier preventing further improvement of ensemble structures.

A useful analogy to density misfit barrier traps is the mechanical over-center, such as a pair of locking pliers. When the pliers are open there is no strain, when they are locked the strain is high, but the highest strain of all occurs just as the pliers begin to open, and this keeps the tool locked in the high-strain clamping state. Because current practice of conformer assignment tends to be random, about half of them are expected to have strained connectivity to the rest of the molecule (locked pliers). They are jammed in place, and cannot relieve this strain by minimization alone.

Prior to this work, the only way to tell if an ensemble model was trapped in this type of local minimum was trial-and-error: swap the altlocs of some atoms and re-refine (§3.4). If the geometry improves, then the starting structure was trapped. That is, if a nudged ball rolls downhill, then it was not yet at the bottom. This process is unfortunately neither efficient nor ergodic, and the purpose of this work was to inspire a better approach.

### 1.3 Alternate locations and correlated motions

Addressing the issue of strained local minima in refinement would open up new areas of investigation in protein dynamics and drug discovery because the models will be more realistic and accurate. Correctly identifying coordinated motions is equivalent to accurate conformer assignments to alternate locations, not just in side-chains, but loops, whole domains and even coupled shifts within active sites, all of which are currently recalcitrant to direct observation. This prediction is based on the assumption that the members of the true, underlying ensemble of conformations that add up to the observed density are each, individually, in good agreement with expected chemical geometry. Here we assert that a low-strain model that fits observed density must exist.

A remarkable finding of this work is that once this lowest-strain ensemble is found, any deviation from it is detectable; either by demonstrably worse geometry or by statistically significant difference-density features. That is, even if this “real” pose were only slightly chemically better than all the incorrect poses, it would still be possible to roll downhill into it, were it not for density misfit barrier traps. This suggests that arriving at the true underlying ensemble for the whole molecule may not be merely sufficient, but necessary to explain crystallographic data to within experimental error. Although proteins are often viewed as flexible structures, they are far more well-organized than other polymers of the same size, and this is likely required for many of their functions. It is also well-known that long-range communication must play a role in regulation and allostery (Changeux, 2012, Raman *et al*., 2016). Even if only partially successful, deducing coordinated motions is expected to have a profound impact on the study of biological function, especially ligand binding as depicted in Figure 2.

The goal of this Challenge was to pave a path to opening these doors. New algorithms were required to specifically target local minima and reliably and consistently generate accurate multi-conformation models. Determining the effectiveness and guiding development of these algorithms is best done with a known ‘correct answer’, or ground truth, which does not currently exist with real-world data, as was done by (Bunkóczi *et al*., 2014). We therefore constructed a synthetic data ground truth that captured the real-world problem of interest, but was not so “realistic” as to be intractable to solve.

## 2. Methods

### 2.1 Creating the Ground Truth Structure

The underlying model defining the target electron density data of this challenge is a 2-conformer ensemble structure generated based on the real-world scorpion toxin 1aho (Smith *et al*., 1997). This protein was selected primarily because it is small (7.6 kDa, 64 amino acid residues), to speed computation, yet contained essential features of proteins, including at least one of each of the 20 most common amino acid residues, and four disulfide bonds. In addition, despite originating from high quality data extending to 0.96 Å resolution, 1aho still had reported R_work_=16.3% between observed and calculated structure factors which was significantly larger than the error of the observations (R_merge_=7.3%, R_σ_=3.8%). A 2-member ensemble was chosen because this is the simplest possible ensemble. Although we found that density misfit barrier traps can occur at any resolution (data not shown), high resolution data was generated to make traps as easy as possible to detect.

Appendix C details the ground-truth model generation. The final model was roughly consistent with the real-world electron density (R_work_/R_free_ = 16.5%/16.8%), but free from chemical strain outliers larger than 3x standard deviations, save three outlier rotamers that were clear in the real density. A central assertion of this work is that although real molecules may contain strain, extreme outliers are not acceptable unless supported by direct experimental evidence.

B factors were optimized in refmac5 (Murshudov *et al*., 2011), because refmac5 has tighter B-factor restraints than phenix.refine (Afonine *et al*., 2012) with default settings. Phenix.fmodel (Liebschner *et al*., 2019) was then used to generate the ground-truth structure factors (F0). Missing reflections from the deposited 1aho data were removed and Gaussian noise derived from deposited standard deviation (SIGF) was added to the F0 structure factors using sftools (Agirre *et al*., 2023) to form F in the distributed data file named refme.mtz.

It is noteworthy here that no attempt was made to simulate non-flat bulk solvent. Solvent modelling will be the subject of a future challenge, not this one. Here we chose to showcase a single major source of systematic error without interference from anything other than expected experimental noise. The bulk solvent parameters were chosen to make it possible for both phenix and refmac5 to re-create the exact ground truth bulk solvent mask internally, and a .fab file of the ground truth solvent was provided for use in SHELXL (Sheldrick, 2015).

Ordered waters were kept and also assigned two alternate locations (altlocs) because clashes with water were anticipated to be helpful in resolving conformer assignment ambiguities. After some initial difficulties with stability of multi-conformer water, it was decided that participants may use the ground-truth water partial structure as a fixed entity, and scripts were provided for restraining the water positions during refinement.

In summary, although the R-factors presented here would have been more “realistic” if we had incorporated a sophisticated solvent model or other sources of non-random error, the solvent and all other systematic errors was purposefully kept simple in order to maximize the possibility of success.

### 2.2 The weighted energy (wE) scoring function

A successful challenge needs a well-defined scoring system, but there are many popular validation scores used in macromolecular crystallography. Appendix B details how these scores were combined, but briefly: All scores were converted to a common scale as statistical energies (as in Equation 1), such as representing rotamer probabilities as the sigma deviate that occurs with that probability. Large statistical energies (>10) were regularized with a plateau (see clip(E) in Appendix B) to prevent tiny changes in a single outlier feature from overwhelming salvageable defects. For example, reducing a 6 σ deviate to 3 σ via a trade-off that increases an outlier already at 20 σ to 20.4 σ was made a favorable compromise by this plateau, despite the difference in squared deviates being larger for the outlier.

The very worst deviate (max(E)) and the average energy (<E>) in each of 11 categories (i.e. bonds, rotamers, clashscore, etc.) were then weighted and summed to form the wE score:

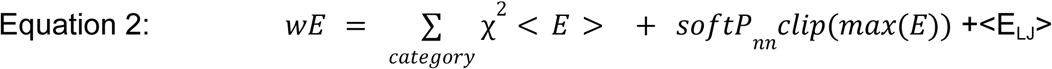

Worst- outlier weights (softP_nn_) reflect the probability that the worst deviate was not noise. I.E. a 6σ deviate is not noise, and gets a weight of 1, and 1σ deviates get a weight of 0. <E_LJ_> is the average Lenard-Jones energy of all non-bond interactions, which can be as low as −1 if all nonbonds are favorable. This was intended to favor well-packed structures. Average energies <E> in all other categories were weighted by the Χ^2^ test, meaning that if the observed deviate distribution is far broader than expected from the geometry σ values, the average energy gets a weight of 1, and if it is far better than expected, it gets a weight of 0. If the observed distribution of deviations is exactly as expected, both weights are 0.5. In addition, clashes and other non-bond interactions were given the much harsher Lenard-Jones penalty than that usually used by refinement programs. As a result of the clipping of energies at 10, the magnitude of the wE score tends to be ∼10x the number of categories with severe outliers, but clashes push it much higher.

### 2.3 Creating the Challenge models

Rather than an all-or-nothing test, the Untangle Challenge was structured as a series of increasingly difficult problems to solve. In each case a model was refined against the same data, refme.mtz, but the models themselves have more and more difficult-to-escape local minima. The Challenge levels are described below:

#### Level 0: Validation

The model called best.pdb was the result of refining the ground truth structure against the ground-truth data in refme.mtz using phenix.refine (Afonine *et al*., 2012) with non-default parameters set to disable occupancy refinement and NQH-flipping, as well as wxc_scale=1.4, because higher and lower values of this weight produced higher wE scores than those of best.pdb: wE=18.2707 with R_free_=3.1%. These are excellent statistics that proved very difficult to beat.

It may seem counter-intuitive that the data in refme.mtz were generated from the starting atoms of this refinement, yet the structure still moves. This is because the refinement that produced the ground truth model had non-default geometry weights, and was refined against real data (see Appendix C). That is, the coordinate file declared to be the “ground truth” came out of a refinement that used the deposited 1aho X-ray data as the X-ray target. Subsequent refinement of the ground truth model against data generated from itself represents the jump in this work from real-world data to synthetic data. It also represents a jump to declaring the statistical energy used by phenix.refine to be the ground truth of chemical energy. This best.pdb was so named because it represents the best wE score than one could reasonably hope to achieve with commonly user-modified refinement parameters.

Remarkably, a wide variety of starting models converged to within a maximum deviate of 0.01 Å from best.pdb, all that was needed was conformer assignments to be on the correct “side” of the electron density, and best.pdb was readily reproduced. This is the model against which subsequent levels of the challenge were judged.

#### Level 1: One thing wrong: Ala39 swap

One model that does not reproduce best.pdb when refined against refme.mtz is otw39.pdb (Figure 4AB). It has one thing wrong (otw) and was generated from best.pdb by swapping conformer assignments of the two C_β_ altlocs of Ala39 and re-refining. Final statistics are notably worse: R_free_=3.4%, wE=63.5, and there were several striking geometry outliers, e.g. the N-C_α_ bond of Ala39 is 11σ too long.

**Figure 4:**
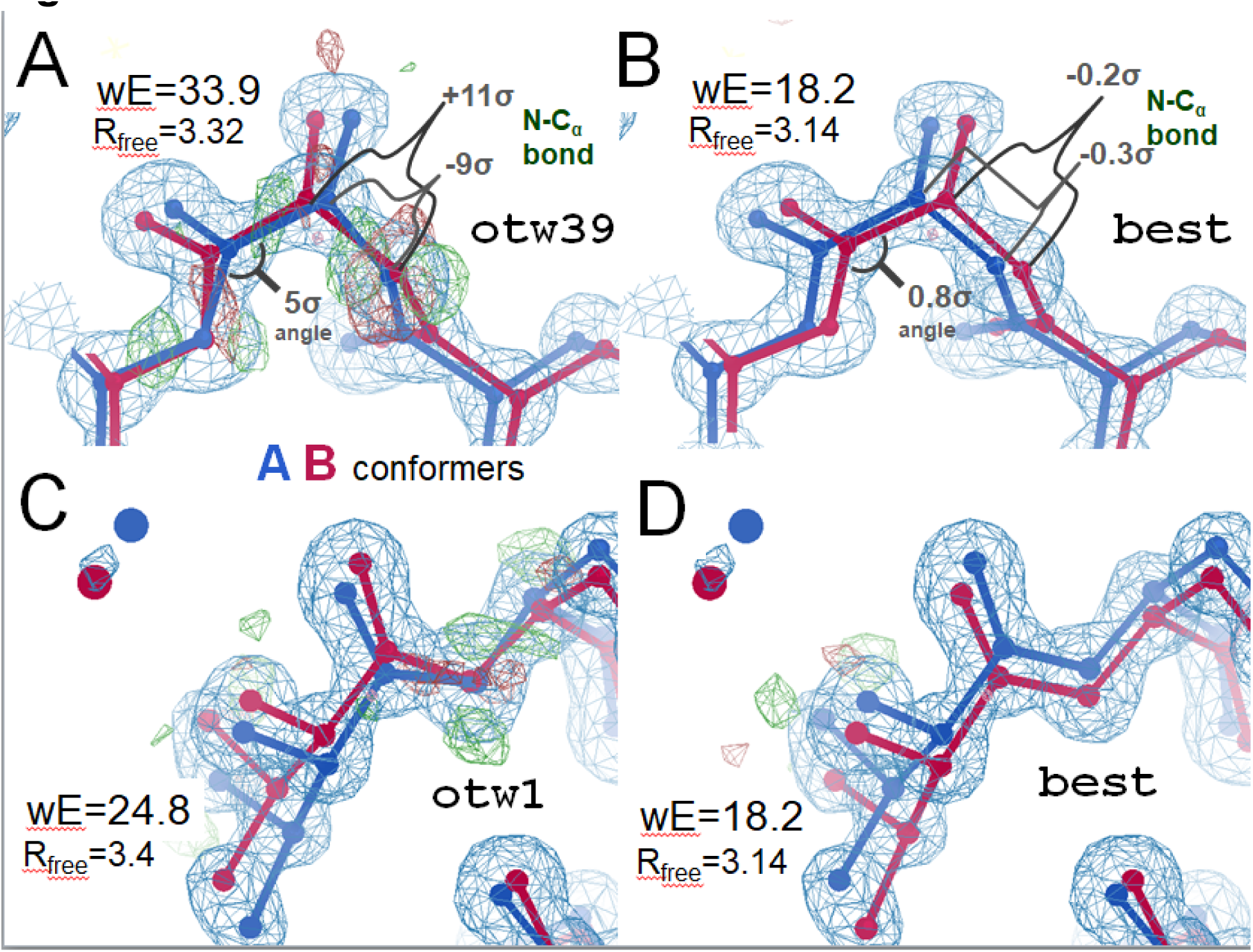
Two examples of small, but stable high-energy local minima due to a density misfit barrier. Top: Level 1 of this Challenge (otw39). (A) large and statistically implausible outliers in bond lengths and angles are denoted in residue Ala39. These strained states remain even after minimization in popular refinement programs such as phenix.refine, refmac5 and SHELXL. However, if either the C_α_ or C_β_ atom has the labels of its two alternate locations swapped, followed by re-refinement, the low-strain ground-truth structure on the right is recovered (B). This trap can also be released by applying the weight-snap maneuver (§3.2). Bottom: (C) Level 2 (otw1), where the entire Val1 residue is swapped relative to the ground truth. This trap is recalcitrant to the weight-snap maneuver because the geometry strain at the cross-over point is too small. It can be released (D) by the swap-and-rerefine maneuver (§3.4).

#### Level 2: One thing wrong: Val1 swap

Similar to Level 1, Level 2 of the Challenge was generated by swapping a single atom and re-refining, this time the C_β_ atom in Val1 (Figure 4CD). The resulting model, otw1.pdb, had poorer stats: Rfree=3.4%, wE=24.8 arising mainly from a Molprobity (Williams *et al*., 2018) clash between Val1 and a nearby water S56. The ground truth has no clashes. Level 2 is more difficult than Level 1 because Level 1 can be solved by adjusting the refinement geometry weight (see §3.2), which is a commonly user-modified parameter. The trap in otw1.pdb, however, was recalcitrant to the weight-snap maneuver.

#### Level 3: Lots of things wrong

The model lotswrong.pdb was trapped in a manifold local minimum similar to the one-thing-wrong levels. The stats: Rfree=4.4% and wE=104 were much higher because instead of just one group of atoms to swap, there were 129 atoms in 75 residues (43 protein, 32 water) that needed swapping. This model had six clashes, 6-sigma bond and angle deviates, C_β_ deviations, and many other problems. Despite the highly disagreeable geometry, this structure remains trapped in all popular refinement packages. But, simply swapping the right conformer letters and re-refining recovered best.pdb. This level is hard because brute-force exploration of all possible combinations of 129 swaps is not a tractable approach.

#### Level 4: Anisotropic B factors

All atoms in the ground truth of this Challenge have isotropic B factors, but two conformers. It is common practice to model atoms with alternate locations (altlocs) sufficiently close together as a single position with an anisotropic B factor. This has the advantage of reducing the number of bonds, angles and other parameters to fit. Moreover, the fused alternative conformers are intrinsically untangled, like the pincer maneuver (§3.5). Nevertheless, the statistics (R_free_=6.6%, wE=59.8) are worse than all but the worst fully 2-conformer models. This is because bonds that connect a 2-conformer to a 1-conformer section of the molecule are always more distorted than they are in a fully single-conformer or ensemble model. The challenge of this Level was to use the conventional anisotropic B approach to match or improve upon the statistics of the wholly 2-conformer ground truth.

#### Level 5: Start with a manually-built model

The starting model for Level 5, manual_built.pdb, was generated through iterative rounds of refinement starting with the qFit model described in Level 6 below. This model has generally good stats, but nevertheless contained a large number of clashes and non-bond interactions, large bond, angle and torsion deviations, a twisted peptide and some bad C_β_ deviations led to wE=93.4 with R_free_=4.56%. These defects were not intentionally introduced, but rather the result of normal, contemporary model-building practices. The low R_free_ and high wE demonstrate how the model can be “shrink-wrapped” into density when the solvent region is accurately modeled (flat in this case) and generates no phase errors. The primary purpose of this Level is to act as a jumping-off point for expert model builders to try their hand at identifying and escaping these local minima using the tools of their choice.

#### Level 6: Start with qFit

One of the few programs written especially for multi-conformer model building is qFit (Wankowicz *et al*., 2024). The model qfit_best.pdb, with R_free_=9.3%, and wE=97.3, was generated from the phenix.autobuild model from Level 7 below using three rounds of qfit 3.0, followed by back-and forth refinement with phenix.refine, refmac5 to decorate hydrogen in the same way as best.pdb and add the OXT, then phenix.refine again with default settings. It contained 28 single-conformer, 29 two-conformer, 20 three-conformer protein residues, and 77 single-conformer waters. This level of the Challenge was designed to allow participants to start with the current state-of-the art in automated multi-conformer model building.

#### Level 7: Start with phenix.autobuild model

The output of phenix.autobuild (Terwilliger *et al*., 2008), had wE=18.1, but R_free_=18.4%. The difference map also has huge peaks (36 σ). This is because phenix.autobuild only builds one conformer. The purpose of this level was to use modern automated model building as a starting point.

#### Level 8: Start with traditional ensemble refinement

Ensemble refinement is a natural choice when you are looking for an ensemble, and phenix.ensemble_refine (Burnley *et al*., 2012), starting with the deposited structure, was used with default parameters to generate phenix_ensemble.pdb. Final statistics: R_free_=9.4% and wE=109.7 are similar but slightly worse than those from qFit. It is also complicated by the fact that it contains 200 models: the more members an ensemble has, the more tangled it can become. Selectively reducing this number to two might be a viable approach to this Level of the Challenge.

#### Level 9: Long-range traps

Density misfit barrier traps become particularly problematic when they are no longer localized, but large regions of the model are swapped in A-B conformation assignments relative to the ground truth. The model longrangetraps.pdb was created using rectified simulated annealing (§3.3), where any atoms knocked out of density or into bad geometry by a simulated annealing (SA) run were restored to their original positions, and the SA repeated. Temperatures ranged from 2000K to 8000K in 27 steps and were combined with 20 random number seeds and weight-snaps (§3.2). The result had very good stats (wE=21.4, R_free_=3.48%), but still not as good as best.pdb (wE=18.2, R_free_=3.14%). Depicted in Figure 5, this model had little localized strain, and the atomic positions are very close to those of best.pdb, except for their A-B conformer assignments. This level demonstrates that not all density barrier traps are small and localized, and that once high-strain traps have been released, what remains are large numbers of atoms that must all be swapped at once to escape these long-range local minima.

**Figure 5:**
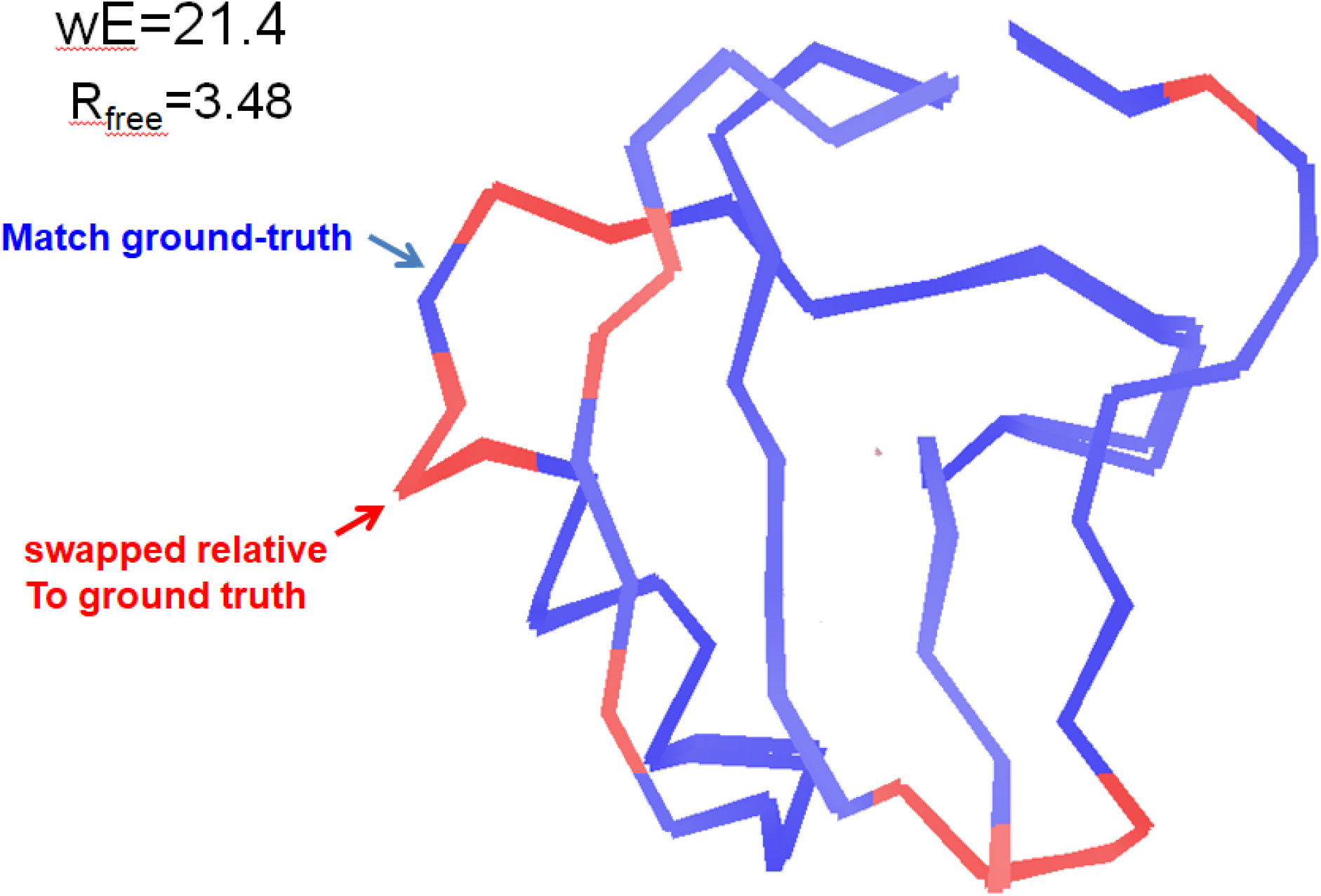
Level 9 Long-range traps: was constructed by applying the rectified simulated annealing algorithm (§3.3) to the structure from Level 3: lots of things wrong. The traps in levels 1-3 are high-strain and relatively easy to identify. Here the overall strain and R_free_ are both low, but not as low as best.pdb. Large and contiguous sections of the molecule match the ground truth, but other regions are flipped relative to it. Unlike the more localized traps in Levels 1-3, the borders between these regions have little strain. Weight snaps (§3.2) or single-atom swap-and-rerefine (§3.4) did not release any of these traps. Only by identifying and flipping large groups of atoms simultaneously can these long-range traps be released. The errors in this model correspond to incorrect assignment of “windshield wiper” vs “jumping jacks” modes of molecular motion.

#### Level 10: Recover the Ground Truth

This was the real Challenge: devise a way to arrive at the global minimum, where geometry energy and density fit are simultaneously optimized. A central thesis of this work was that although many models may have good density fit and reasonable geometry, the wE score, and, to some extent, R_free_ betray them as being trapped in local minima. The global minimum appears to be identifiable, but an algorithmic path to arriving at it stating with density data alone appears elusive. This level of the Challenge is meant to be open to all approaches and ideas, without requiring a specific starting model. The only thing deemed to be cheating was using best.pdb as a guide.

#### Level 11: Alternative hypothesis

The second major goal of this Challenge was assessing the uniqueness of the global minimum. This was because a critical factor in obtaining the real-world underlying ensemble is proving that no alternative hypotheses is plausible. A central postulate of this Challenge is that the members of the true, underlying ensemble do not contain highly strained and statistically implausible outliers of chemical strain, unless the density data clearly support them. The Challenge at this Level was to build a model into refme.mtz that represents a potentially functionally relevant alternative to the ensemble represented by best.pdb, and also has as-good or better R factors and validation statistics. That is, find a model that does “jumping jacks” instead of “windshield wipers”, and scores better than best.pdb.

## 3. Results and Discussion

After creating the Challenge Levels described above we tried various methods to solve them. Many approaches were ineffective, but others proved useful. Of particular note were techniques we are calling: the weight snap maneuver, rectified simulated annealing, swap-and-rerefine, and the pincer maneuver. The Phenix and ROPE groups created new software. In addition, several algorithms were developed by third parties in response to the Challenge, which we will describe briefly below.

### 3.1 Alphafold2

It is now very common to start a structure solution with a predicted model, but we found this was not an effective strategy for avoiding tangles. Structure prediction programs such as Alphafold (Jumper *et al*., 2021) and RosettaFold (Baek *et al*., 2021) do not yet produce altlocs, but do provide a list of top-scoring models. The top five models from Alphafold2 prediction of the sequence of 1aho were paired up, two at a time, and refined against the Challenge data as a 2-conformer system in the context of waters fixed at their ground-truth positions. The best result after refining all possible combinations had wE=129.5, R_free_=13.54%, which was the worst geometry of all the models presented here. The best of all five single-conformer predictions had wE=155, which we note was unrelaxed. A round of phenix.geometry_minimization brought wE=7.97, but severe disagreement with density (R_free_=56%).

The biggest problems with the Alphafold2 predictions were that the C terminus was placed 8 Å away from its correct location, and none of the strained rotamers were predicted. This placed all the Alphafold2 models outside the convergence radius of refinement. To simulate the most favorable manual rebuilding possible, we applied an iterative “cheating” maneuver, where the single most mis-placed atom was moved and restrained to its ground-truth position for further refinement. This was then repeated for the next most mis-placed atom, and so on until the ground truth was recovered.

Of all pairwise starting points, arriving at an in-density model (R_free_<4%) took a minimum of 5 rounds of “cheating” moves of more than 1.5 Å. This was started with Alphafold2 predictions ranking 1 and 3, and then moving His64, Asp9, and Cys63 by into density, arriving at wE=51.95. Recovering the ground truth took a total of 14 rounds of cheating moves, with wE dropping below 18.5 at round 12 with two small moves (O of Gly4 and Oγ of Ser40) less than 0.3 Å left to arrive at wE=18.2. Without knowing which of the two alternative positions in the ground truth to use for each round, performing this task by exhaustive search would have been a formidable combinatorial search without prior knowledge of the ground truth.

### 3.2 Refinement weight snap

Many types of density misfit barrier traps could be resolved with a simple 3-step procedure referred to here as a “weight snap maneuver”. The relative weight given to geometry vs X-ray residuals (wxc_scale in phenix or “weight matrix” in refmac5) was given a very high value, followed by a very low value, and then the default value again. This temporary high prioritization of either residual in turn can overcome the stalemate between them and resolve the trap. Level 1 is such a case, Level 2 is not, but as a general tool snapping or sometimes ramping the weight was found to almost always improve ensemble models, and no cases were found where it degraded them.

### 3.3 Rectified simulated annealing (RSA)

Simulated annealing (SA) is a classic approach to escaping local minima, but for the tangling problem it tends to fall into about as many traps as it escapes. These transitions into higher-energy states, however, can be identified and reversed because the force field can be evaluated on an atom-by-atom basis. That is, for a single atom, all the bonds, angles, torsions, clashes, and other energy terms that involve the atom of interest can be evaluated before and after a standard SA run. If the aftermath is significantly worse, then the atom can simply be restored to its original position and the entire structure re-refined. This one-way valve approach is analogous to rectification with a diode in an electronic circuit, or a back-cross in genetics, where wild-type sequence is restored everywhere except where necessary to maintain the desired phenotype. Except here the sequence being manipulated is a series of conformer letters.

### 3.4 Swap-and-rerefine

Perhaps the most potent way found to escape a density-misfit barrier trap was to leap over them. A trial atom with two alternate locations can simply have those conformation assignments swapped, and then the entire structure re-refined. With only 582 non-H atoms in the ground-truth structure, swapping all of them one at a time was straightforward on a modestly-sized computer cluster. The problem with this approach arises when one considers swapping more than one atom at a time. Exhaustively swapping all 169,053 possible pairs of atoms was tractable with high-performance computing, but swapping triplets, even for this small protein, was only partially explored. Also, in cases with more than two conformers all possible permutations of those letters must be checked, vastly increasing computational cost. Nevertheless, applying trial single-atom conformer swaps was found to be an effective and straightforward way to improve any multi-conformer model, and a script for performing such swaps was provided with the Challenge data.

### 3.5 Pincer Maneuver

Another way to escape a density misfit barrier trap was to position both altlocs at the exact same location: the centroid of the density, or halfway point between them. It is best to restrain them to stay at this location in an initial refinement run and let the rest of the structure relax in that context. Essentially, the two alternate locations are “pinched” at the top of the “hill” in terms of the density misfit energy. This gives the geometry term the maximum chance of pulling the model down the correct side of the density once the pincer restraint is released. Overall, however, we found pincer maneuvers to be somewhat less effective than swap-and-rerefine.

### 3.6 Phenix

Within tthe Phenix Suite, Terwilliger implemented the wE score in *phenix.holton_geometry_validation*, and developed a general procedure called *phenix.create_alt_conf* that uses wE as a target function optimizing the arrangement of conformers in an ensemble model. It systematically varies the arrangement of conformers, identifies patterns in the scores of the resulting models, and attempts to generate new arrangements with low wE scores. Combined with a final scan of wxc_scale values to minimize wE, this procedure was successful in generating models with wE scores and R_free_ values very similar to those obtained for best.pdb at levels 1-5 and 7-9 (Table 2, the other levels were not attempted). Note that for levels 0-2 the arrangement of conformers was identical to best.pdb, but for the higher levels the arrangements found were quite different.

**Table 1:** Summary of weighted geometry energy (wE) and R factors for various approaches to untangle Challenge models.

| wE | R<sub>work</sub>(%) | R<sub>free</sub>(%) | method |
| --- | --- | --- | --- |
| 129.487 | 12.91 | 13.54 | AlphaFold2 - best pair |
| 109.6 | 8.87 | 9.44 | phenix.ensemble_refine |
| 97.3 | 8.84 | 9.27 | qFit default |
| 91.7 | 3.64 | 4.27 | Manual built |
| 80.5 | 4.31 | 4.75 | L3: lotswrong |
| 54.8 | 9.47 | 10.4 | Amber24 |
| 54.6 | 5.87 | 6.57 | L4: aniso B with few splits |
| 33.5 | 2.94 | 3.32 | L1: One thing wrong: Ala39 |
| 23.5 | 2.95 | 3.31 | L2: One thing wrong: Val1 |
| 21.4 | 3.12 | 3.48 | L9: Rectified Simulated Annealing |
| 19.4 | 3.72 | 4.64 | ROPE GUI (by hand) |
| 18.1 | 17.9 | 18.4 | L7: phenix.autobuild |
| 18.2 | 2.79 | 3.14 | L0: best.pdb - refined ground truth |
| 17.6 | 3.18 | 3.71 | phenix.create_alt_conf |

**Table 2:**
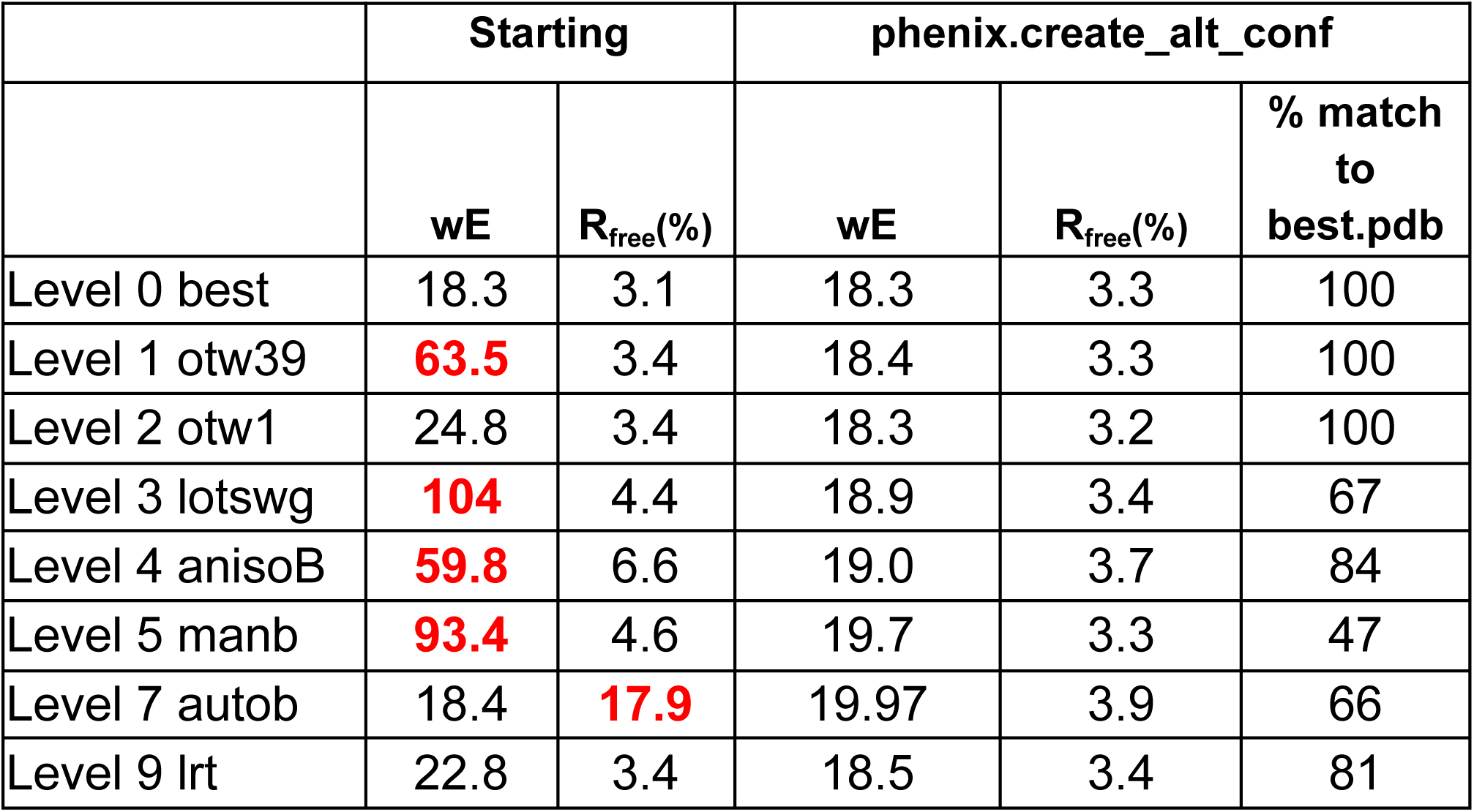
Phenix create_alt_conf applied to Challenge models. Each starting model was analyzed with default parameters except the number of models generated in each cycle was 640, H atoms that are not recognized by phenix.refine were ignored, the H-bond donor-acceptor distance used in generating the Challenge was used (const_shrink_donor_acceptor=0.6), and water sites were chosen automatically from a list of all the actual water molecule sites (fixed_solvent_model=perfect_water.pdb). At the conclusion of each run of create_alt_conf, the solution with the lowest wE was re-refined with phenix.refine using anisotropic B factors and a range of refinement weights (wc) ranging from 0.5 to 8 and choosing the one with the lowest resulting wE value. The Free R values and wE scores after simple refinement and after application of this procedure using create_alt_conf are listed.

|  | Starting |  | phenix.create_alt_conf |  |  |
| --- | --- | --- | --- | --- | --- |
|  | wE | R<sub>free</sub>(%) | wE | R<sub>free</sub>(%) | % match to best.pdb |
| Level 0 best | 18.3 | 3.1 | 18.3 | 3.3 | 100 |
| Level 1 otw39 | **63.5** | 3.4 | 18.4 | 3.3 | 100 |
| Level 2 otw1 | 24.8 | 3.4 | 18.3 | 3.2 | 100 |
| Level 3 lotswg | **104** | 4.4 | 18.9 | 3.4 | 67 |
| Level 4 anisoB | **59.8** | 6.6 | 19.0 | 3.7 | 84 |
| Level 5 manb | **93.4** | 4.6 | 19.7 | 3.3 | 47 |
| Level 7 autob | 18.4 | **17.9** | 19.97 | 3.9 | 66 |
| Level 9 lrt | 22.8 | 3.4 | 18.5 | 3.4 | 81 |

These results demonstrate that there is a wide and shallow basin of tangled models that explain the density well (low R_free_), but are “shrink-wrapped” into it. Higher wE score generally differentiated between these local minima and the ground truth topology. In other cases careful adjusting of refinement parameters can meet or surpass the ground truth wE, but then the difference density is telling.

For example, in Figure 6 we show one create_alt_conf result with wE score better than that of best.pdb, but difference density features indicate there is something wrong. Further refinement of this model with the same settings used for best.pdb yielded wE=58.4 and R_free_=3.4%. This result demonstrated how R factors and other global metrics are weak indicators of density fit. We plan to develop better density scores in the future.

**Figure 6:**
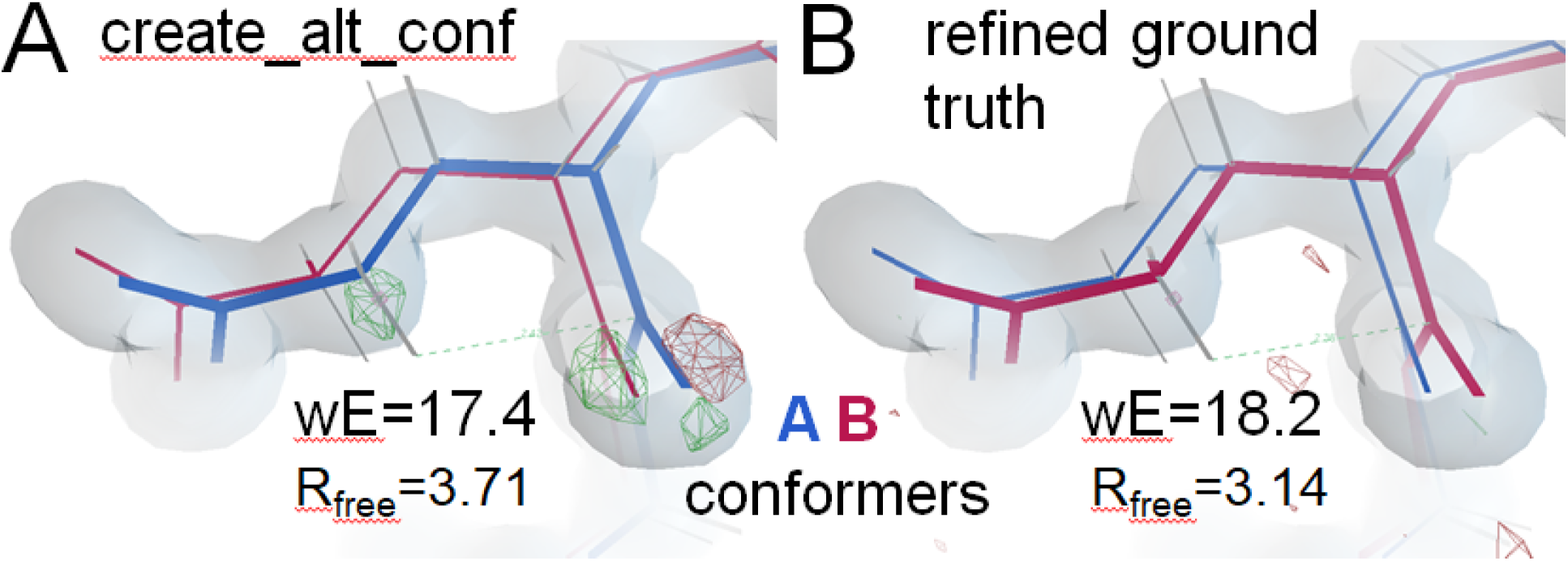
Example of a structure (A) with better wE than best.pdb (B) derived from phenix.create_alt_conf. The improvement in wE score is due mostly to better average geometry deviation energy. Two worst-outliers also improved: non-bond interaction: Hγ_2_ vs carbonyl C and torsion χ2, both of Glu32. This residue is one of the three *bona-fide* outliers observed in the real 1aho structure that was kept in the ground truth of this Challenge. Increasing the geometry weight improved the geometry, but pulled the atoms visibly out of alignment with the density. The mF_sim_-DF_c_ (red and green nets) was contoured at 5σ in A, and at 3σ in B. The change in R_free_ is small, but at this low R level subtle mis-matches are readily apparent as difference features up to 11σ tall (not shown). Employing anisotropic B factors suppresses this difference density, making it much more difficult to identify the ground-truth configuration.

### 3.7 RoPE GUI

Ginn used the RoPE framework (Ginn, 2023) to create what is perhaps the first GUI to display alternate conformers (altlocs) of atoms with different colours by default; the two altlocs are drawn in blue (A) and yellow (B) (Figure 7). Starting with lotswrong.pdb (Level 3), which has every atom in two altlocs, four geometry scores were evaluated for each covalent bond. The starting-model (cognate) assignments were referred to as AA and BB, and bonds that would be formed after swapping altloc assignments (non-cognate) were AB and BA. For each of these combinations, due to the different altlocs of A and B, there are four distinct bonds which can be drawn (represented as rope in the GUI). Bonds between AA or BB were drawn with a natural hemp rope colour, and bonds between crossed altlocs (AB or BA) were drawn as a gradient between the two respective colours (a blue-yellow gradient or yellow-blue gradient). The geometry scores were used to modulate the transparency of the bond. Bad geometry bonds were made more transparent. Therefore, a correctly assigned region would predominantly have a normal rope colour at full opacity. Regions with incorrectly assigned alternate conformers tended to have partially opaque cross bonds (Figure 7A). The user may click on an atom to swap the altloc assignment of the two atoms as well as all other atoms downstream of it (toward the bottom of Fig 7). This was not propagated down disulphide bonds or circular amino acids, to avoid infinite loops. The scores were recalculated and the graphics updated to allow the user to decide to accept the alteration or reverse it again based on the colour improvement. This was built into a cyclical set of refinement rounds, each of which involved alternate structure refinement and untangling of chains in the GUI (Figure 7B). Refining with a high X-ray residual weight was found to enhance the contrast of geometry swaps suggested by the colour scheme, and the result of swapping the altlocs is shown in the case of three regions of the structure (Figure 7C). We were able to achieve wE=19.4 using this interface.

**Figure 7:**
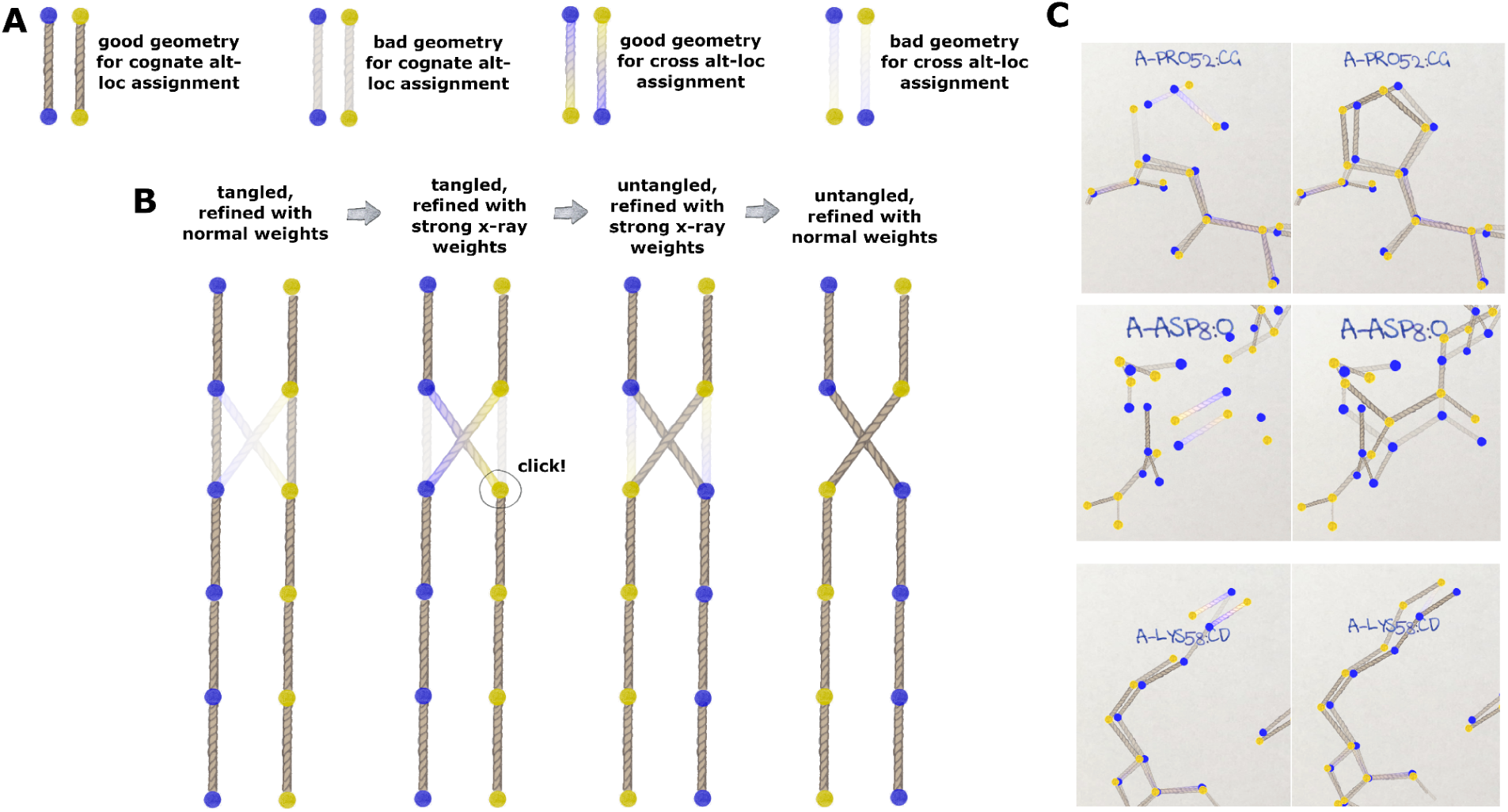
Behaviour of RoPE GUI for untangling crossed conformations: (A) demonstration of colour changes between altloc combinations (either colour gradient or single hemp colour) and geometry scores (bad scores more transparent; good scores more opaque). (B) workflow of operations: 1st: the normally-refined tangled structure shows hints of a conformer swap in the second bond from the top; 2nd: using the first refinement of the “weight snap” method with a high x-ray residual term strengthens the hint of the suggested conformer swap; 3rd: user can click on the atom to be flipped, which propagates downstream; 4th: further refinement with normal weights improves the cognate geometry. (C) examples of tangled regions of lotswrong.pdb refined with a high X-ray weight on the left and after flipping the altloc assignment on the right.

These manipulations were blind to the ground truth, and in post-analysis, 23 atoms in 5 residues remained swapped relative to the ground truth: most of Leu51-His54, O/Nε of Gln37. OH of Ser40, and the carbonyl O of Cys46. This last carbonyl trap caused the most residual strain, but not to itself. Swapping this carbonyl O makes the C-C_α_-C_β_ bond angle distort, making it difficult to see which atom needed to swap.

### 3.8 Third-party contributions

Since the initial announcement of this Challenge several groups have reported to us on their efforts to solve it. Some are authors of the present work, but other noteworthy contributions are briefly described here. One such case was (Mandaiya & Elser, 2025) used a Divide and Concur Framework and a Reflect-Reflect-Relax search algorithm to successfully untangle the otw39.pdb and otw1.pdb, respectively.

David Case used Amber24 (Case *et al*., 2025) to conduct molecular dynamics simulations in the context of electron density restraints using an annealing-based protocol shown to substantially improve model statistics (Mikhailovskii *et al*., 2024). The result: wE=54.8 and R_free_=10.4% is a substantial improvement over manual building attempts, except for RoPE (§3.7).

Dale Tronrud proposed a “variable labels” approach (Tronrud, 2023) where atomic positions are given not just one name, but a probability distribution of names. These probabilities could then be differentiated against the energy terms and refined like any other parameter. This novel approach would avoid the intermediate chain-crossing states and their poor density fit that prevents conventional minimizers from escaping density misfit barrier traps. This algorithm, however, has yet to be implemented in an extant refinement program.

Spencer Passmore has claimed a solution to levels 10 and 11 using a Linear Optimizer, an algorithm often used to approach the classic Travelling Salesman Problem. This approach https://github.com/Phoelionix/Untangler/tree/more_than_2 will be written up in a future report.

## 4.0 Conclusions

This work demonstrates a formidable, yet previously underappreciated, barrier to creating accurate macromolecular models from density data. These deep and non-obvious local minima are resistant to standard refinement procedures, even in the highly favorable case studied here: a small protein with high-resolution data, excellent geometry and a deliberately simplified solvent model. These conformer-assignment-based density misfit barrier traps arise from the topology of the starting model in the refinement energy landscape, where density agreement and geometry restraints can jointly stabilize incorrect solutions. That such failures occur under these idealized conditions underscores how challenging the problem is with real data.

Density misfit barrier traps contribute to high R factors of macromolecular crystallography via the weighting scheme that all refinement programs must use to balance the geometry and density fit energy terms. Single-conformer models do not fit ensemble density well, and ensemble models get tangled and strained. The poor geometry in both cases forces de-prioritization of the density fit. Other systematic errors, arising from non-flat bulk solvent, model/data incompleteness, anharmonic motion, radiation damage, and limitations of current force fields, are undoubtedly important as well, but the results presented here show that even when all these factors are controlled or eliminated, density misfit barrier traps alone are sufficient to prevent conventional approaches from reaching the global minimum. These traps alone are sufficient to explain why the tug-o-war between geometry restraints and density fit persists. These traps also explain why ensemble models thus far have never been able to reach arbitrarily low R_work_ values that would be expected from optimization with a low observations-to-parameters ratio.

A central contribution of this work is the construction of a well-defined ground truth dataset that has statistically plausible and excellent agreement with both the density data and prior knowledge of chemical geometry. The availability of a “right answer” fundamentally changed how new algorithms can be evaluated: failure, success, and progress in between are no longer matters of interpretation, aesthetics, or competing opinions on sources of error, but can be judged directly by proximity to the true solution. This allowed algorithm developers to distinguish methods that genuinely approached the global minimum from those that merely traded one defect for another. We argue that such ground-truth benchmarks are essential for meaningful progress on ensemble refinement.

The weighted energy (wE) score introduced here provided a practical means of ranking models by putting a variety of validation methods onto comparable footing, allowing trade-offs to be judged automatically. Several iterations led to the current state of this tool, and we plan to improve upon it in the future by including all outliers instead of just the worst one into the score, modifying clipping plateau function so that it scales better to larger molecules, and testing against a larger pool of high-quality structures.

Finally, and perhaps most encouragingly, this Challenge has already achieved one of its primary goals: it has motivated the development of new code and new ideas. Multiple independent groups have implemented novel algorithms, search strategies, and user interfaces specifically aimed at escaping density misfit barrier traps, several of which succeed where previous automated tools did not. This collective response strongly suggests that the limitations exposed here are not inherent to macromolecular data, but to the current generation of algorithms.

In summary, ensemble refinement remains challenging, and no single method yet provides a universal solution. However, by isolating a specific failure mode, providing a controlled benchmark with a known answer, and stimulating concrete algorithmic innovation, this work laid a foundation for systematic progress. As these methods mature, we anticipate that improved handling of ensemble topology will become an important part of closing the long-standing gap between experimental data quality and macromolecular model accuracy. Simultaneously, these improved models are expected to better capture the underlying concerted motions of the molecules themselves, which is expected to lead to new functional insights and improved predictive power in drug discovery.

## Acknowledgements

We would like to thank Dr. Tom Peat for his contributions to the early stages of Challenge development, and the manual-built model, and Drs. Dorothee Liebschner, and Nigel Moriarty for their advice with phenix.refine. This work was supported by grants from the National Institutes of Health (R35 GM158447, P30 GM124169, P30 GM133894, P50 AI150476, and U54 AI170792), and the US Department of Energy under contract No. DE-AC02-05CH11231 at Lawrence Berkeley National Laboratory and DE-AC02-76SF00515 at SLAC National Accelerator Laboratory. P.V.A. acknowledges funding from the National Institutes of Health (grants R01GM071939, P01GM063210, and R24GM141254), as well as support from the Phenix Industrial Consortium and the US Department of Energy under Contract No. DE-AC02-05CH11231. Data files are available at: http://bl831.als.lbl.gov/∼jamesh/challenge/twoconf/ and https://github.com/jmholton/UnTangle.

## Appendix A Details of how Figure 1 was calculated

In order to demonstrate realistic experimental errors with a defined ground truth a full-scale simulation of a diffraction data collection experiment was conducted using nanoBragg (Wittwer *et al*., 2024, Mendez *et al*., 2024, Sauter *et al*., 2020, Lyubimov *et al*., 2016, Kirian *et al*., 2011). The simulated detector was a Dectris Pilatus 6M with a silicon sensor layer 450 micron deep and square, 172 um wide pixels. This detector was set to be 640.2 mm from the sample, which put spots of resolution 3.1 Å on the edge of the detector at X-ray wavelength 0.977794 Å. X-ray flux was set to 10^12^ photons/s into a 30 micron diameter beam size with divergence 0.1 deg in both directions and spectral dispersion of 0.2%. This was to simulate unit cell dispersion (Nave, 2014) in the protein crystal as well as the bandwidth of the X-ray beam. Exposure time was set to 0.1 s with 1.0° rotation per image for 360 images. The crystal orientation was randomly chosen to have Arndt-Wonacott misseting angles of −63,−19,-22°. Photon-counting shot noise was simulated assuming the expected detector gain of 1.0. One of the images is shown here, displayed in ADXV (Arvai, 2012):

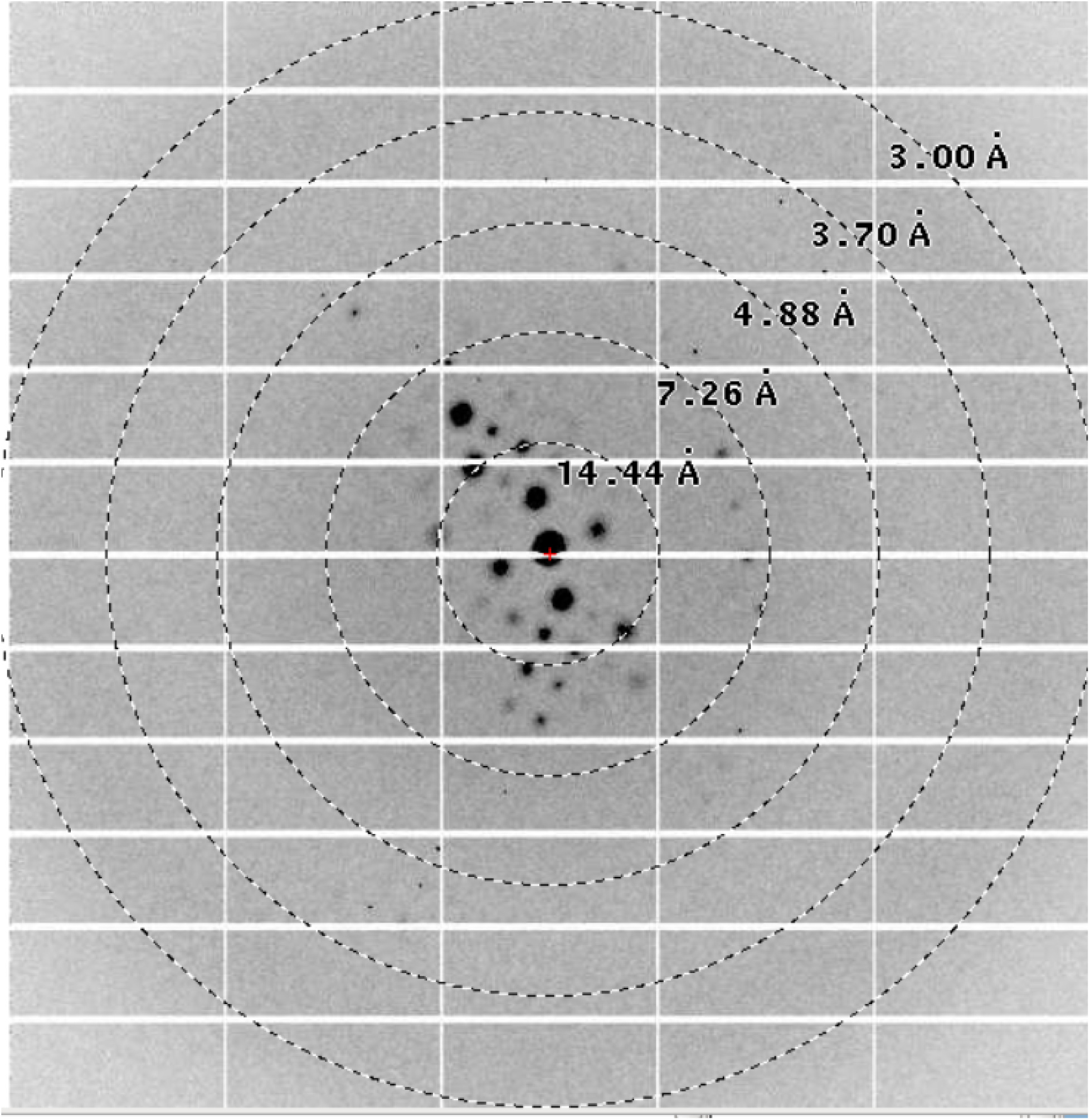

To generate spots with diffuse foothills around the peaks, the Bragg spots were summed over three sub-crystal simulations with the same orientation, but mosaic domain sizes 200, 15, and 5 nm with total scattering contributions scaled up to spheres with diameters 21, 17 and 15 microns, respectively. The crystal was made to have a mosaic spread of 0.2°, which was performed by generating 50 randomly-oriented mosaic domains. The Φ rotation was similarly broken up into 5 steps per image, pixels were over-sampled 4×4, and detector parallax was modeled by dividing the 450 micron thick sensor into 5 layers, each absorbing and transmitting into the next. This level of discretization was found to be sufficient for good data processing statistics without significantly impacting simulation speed.

Background from non-crystalline material was set to be that of a 25 micron thick layer of liquid water, plus 25 mm of air between the collimator and beamstop. In addition, centrosymmetric diffuse scattering consistent with the Wilson B factor of 178 given to the Bragg data was included, as was expected Compton scattering.

Detector calibration errors were simulated by generating Gaussian-random pixel noise and smoothing this noise with a 2D Gaussian kernel with σ-width of 5 pixels. This mask was then scaled to have a root-mean-square (RMS) variation of 3%, and the same calibration error mask was used for all images. Bad pixels were introduced at 600 random locations, in addition to the usual blank areas of this detector type.

Notable sources of error that were not simulated were: sample self-absorption, beam flicker, shutter jitter, sample vibration, dust on the detector window, and other sources of relative error beyond detector calibration. These were neglected here because at modern beamline facilities they are easily made small enough to be dwarfed by the 3% error given to detector calibration in this simulation. Modern detectors can have calibration errors of 1% or less, so this level of overall fractional error was pessimistic. Radiation damage was not simulated because any plausible model of damage can also be used to correct for it. This is equivalent to assuming that effective dose mitigation strategies were employed, such as averaging over many crystals, or zero-dose extrapolation.

The image data were then processed with XDS (Kabsch, 2010) using default settings, and final statistics supported a resolution cutoff of 3.1 Å:

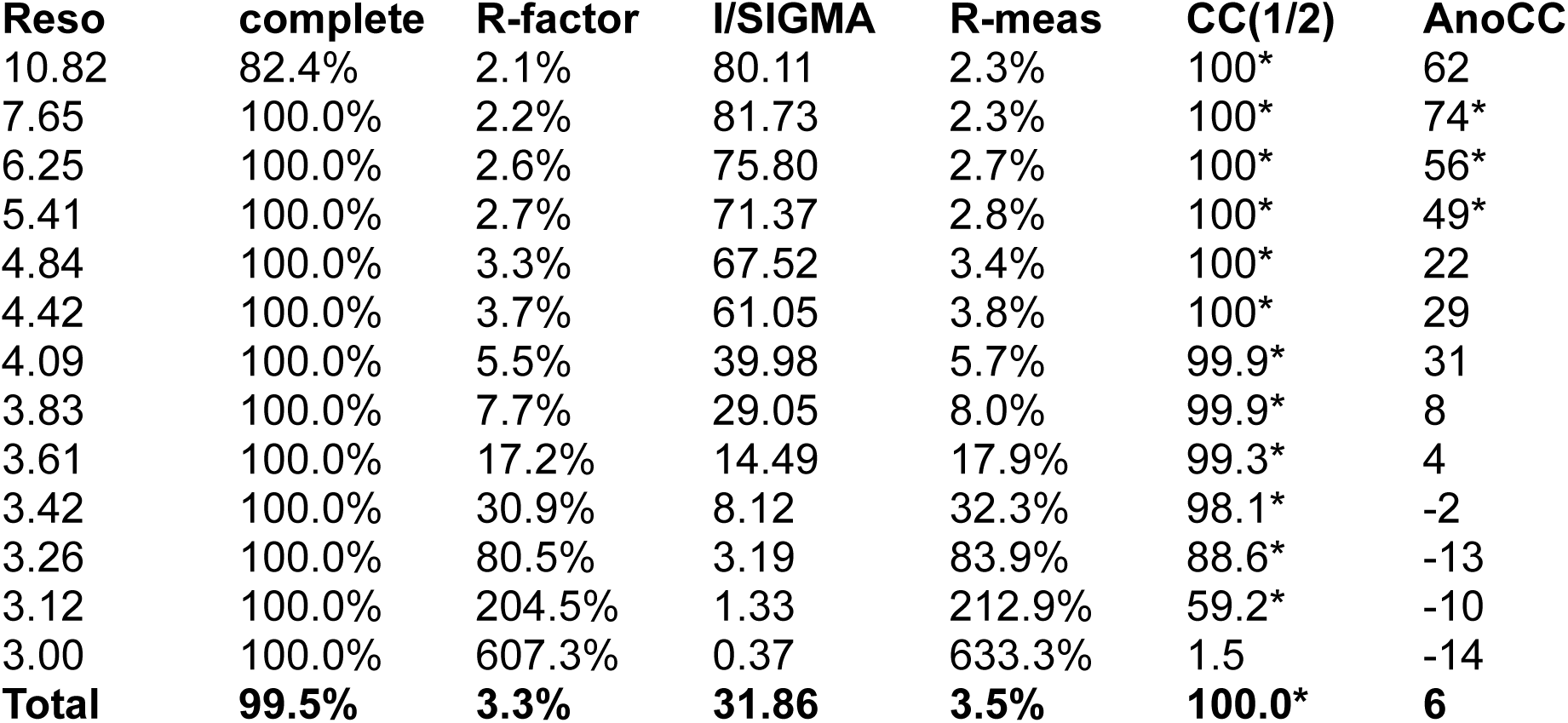

The AIMLESS (Agirre *et al*., 2023) resolution estimate was 3.07 Å.

Post-processing of the merged structure factors was performed to maximize the peak height of the missing hydrogen atom. This was a combination of resolution cutoffs and sharpening and the optimal values found where: sharpening B=-88, refinement data resolution cutoff of 3.556 Å, and difference map resolution cutoff of 4.657 Å. The ground truth model was edited to remove HE1 of His side chains before performing one round of refinement in refmac5.

The ground-truth structure factors were calculated from the best-fit model to the deposited 1aho data obtained to date. This model was derived from a molecular dynamics simulation of 1aho similar to that described in (Cerutti *et al*., 2010), which used a supercell of 2×2×3 unit cells for a total of 48 copies of the protein, 84 acetate ions, 36 ammonium ions and 8760 explicit OPC waters. All non-hydrogen protein atoms and 2297 ordered waters were restrained to crystallographic positions with a weight of 10 kcal/(mol·Å²) for a 100 ns run. Electron density of solvent and other non-protein atoms was averaged over this trajectory. The final frame of the simulation was then refined against the deposited 1aho data in refmac5, using the trajectory-averaged solvent density as a partial structure instead of the default bulk solvent. The hkl data were expanded and re-indexed as 2h,2k,3l to refine the 48-copy structure as a supercell ensemble. After editing the solvent structure, the final R_work_=0.09465 R_free_= 0.11730 were acheived. This rather complex process was used because this model was closer to the observed X-ray data than any other model. It was not, however, used as the ground-truth for this Challenge because high-copy ensemble models like this were found to be unstable in standard refinement programs (not shown).

This 48-copy ensemble model was then energy-minimized in Amber, single-ASU electron density calculated and added to the bulk solvent to form the ground-truth F_true_ for the diffraction image simulation. Crystal disorder was simulated by applying a B factor of 160 to F_true_ before generating spots. The Wilson B factor of F_true_ was 38, so the Wilson B of the structure factors used in the simulation was 178. These B factors were arrived upon by systematically repeating the diffraction image simulation with different B factors and re-processing in XDS until the desired resolution limit of ∼3 Å was obtained.

## Appendix B Details of wE scoring function

A successful challenge needs a well-defined scoring system, but there are many popular validation scores used in macromolecular crystallography, and how to properly balance a gain in one vs a loss in another had to be addressed. For example, bond lengths, angles, torsions, and planar distortions are often restrained using a statistical potential (Eq 1), but Ramachandran, rotamers, and peptide Ω outliers are scored by a probability based on how frequently they are observed in high-resolution structure databases. This was reconciled by converting the probability into its equivalent σ-scaled deviate (Z):

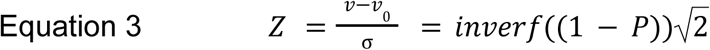

Where P is the frequency of a rotamer or other rare feature seen in databases, and inverf() is the inverse of the Gaussian Error Function, which maps the probability of a Gaussian deviate to the σ-scaled deviation itself. For example, consider a rare rotamer given a score of 0.5% by phenix.rotalyze (Hintze *et al*., 2016). This rotamer is observed as often as one expects to see a 2σ deviate, and inverf(0.995)= 2. The scale factor of 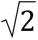 is required because only positive deviates are being considered, making the probability distribution one-sided. This σ-deviate was then fed into Eq 1 to form an energy on the same scale as the other statistical energies.

C_β_ deviations (Lovell *et al*., 2003) were given a statistical potential with σ = 0.05 Å, which was observed to be roughly the root-mean-square variation of C_β_ deviates in low-energy models obtained here.

Non-bonding repulsions usually use a quartic anti-bumping term only (Grosse-Kunstleve *et al*., 2004), and the Richardson clash score (Richardson *et al*., 2018) is a count of interactions that are so close as to be arguably physically impossible. For the wE score, non-bond interactions were replaced with a Lenard-Jones 6-12 potential (Jones, 1924, Hirschfelder *et al*., 1964) commonly used in molecular dynamics that is much more unforgiving of clashes than quartic anti-bumping. Molprobity (Williams *et al*., 2018) clashes were effectively double-counted, as both a clash and a bad non-bond were each given their own energy contribution. Moreover, the number of clashes was also taken as an “energy” to add to the wE score. The ground truth model had no clashes.

The σ of peptide bond Ω deviations was lowered from 5° to 4° because large Ω deviations were found to be common in badly tangled ensemble models (not shown), indicating they are the “weakest link” in the main chain. That is: tangles increase the path traversed by the main chain, forcing something to stretch.

After converting all scores to a statistical energy (E) for all 11 categories: bonds, angles, planes, torsions, chiralities, rotamers, Ramachandran, peptide Ω, C_β_ deviations, non-bonds and clashes, the worst value in each category was weighted by estimating the probability that it occurred by random chance:

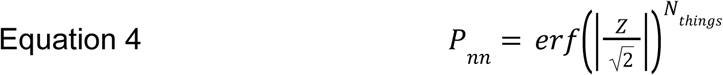

Where erf() is the Gaussian Error Function, or integral over a Gaussian distribution of unit variance. N_things_ is the number of items in the category, such as the total number of bonds in the structure. The number of items in the list affects the probability because the more items are on the list the more likely an extreme value will occur. For example, 4 σ deviates are very rare, expected to occur only once in 6.5×10^7^ events, but if 10,000 events are taken together, the largest deviate will be 4 σ or more 53% of the time.

Taken individually, statistical energies are useful in relating how unlikely a given feature is, but problems were encountered judging a large system collectively. For example, an optimizer that could find a way to reduce forty 1 σ deviates to near 0 σ, but in the process create a single 6 σ deviate, would find the total statistical energy more favorable: 1 x 6^2^ < 40 x 1^2^. But, because a 6σ deviate is expected to be extremely unlikely, and a field of forty 1σ deviates is unsurprising, we devised a weighting scheme to overlay on top of the statistical energy.

The behavior of P_nn_ for large N_things_ was found to be very sharp and numerically unstable, so it was substituted with a softened approximation:

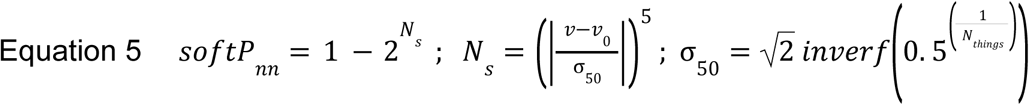

The softP_nn_ function crosses 50% near the same σ-scaled deviation (Z) as P_nn_, but “turns on” more gradually, avoiding sharp dependencies on exact values of parameters, which are themselves uncertain. The resulting probability (softP_nn_) was treated as a weight (w) for the energy (E) of the worst outlier in each category. So, if the worst deviate is very much smaller than expected, it will be given zero weight, and if it is so large as to be extremely unlikely to have happened by chance, it gets a weight of 1.

Now, because there are genuine outliers in the ground truth, it was found that very small changes in the magnitude of the worst outlier were washing out the influence of other categories. To ameliorate this, we applied a regularization that flattens the curve of E once it rises above 10. Specifically:

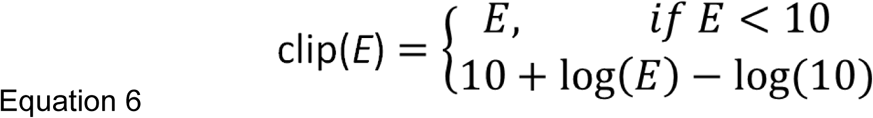

This function continues to rise monotonically as the outlier gets worse, but after passing E=10 it rises much more slowly than E itself. This proved especially useful in balancing clashes and bad non-bonds, as the Lennard-Jones is very unforgiving at short distances.

Finally, the average energy in each category was also weighted by the Χ^2^ test, which conveys the probability or confidence that a given collection of deviates came from a Gaussian distribution. So, if the observed deviations are better than expected given the error bars assigned, the average energy gets weighted to zero, but the more non-Gaussian it behaves, the higher the weight, up to 1.0. The final, overall equation for wE is given as Equation 2. The E_LJ_ shown there is a 12th category for average energies: the signed average of all non-bond L-J energies, which allowed models with “good packing” to score better than models with non-ideal non-bonded distances. The exact break-down of the wE score for the best.pdb model is enumerated in this table:

| category | bond ang | Non-bond | pept $\Omega$ | side $\chi$ | Cβ dev | plane | bond len. | chir | rota | rama | sum |
| --- | --- | --- | --- | --- | --- | --- | --- | --- | --- | --- | --- |
| $N_{things}$ | 1420 | 31 | 126 | 628 | 114 | 1192 | 1046 | 134 | 108 | 124 | 4923 |
| max $\delta/\sigma$ | 1.8 | (2.4) | 2.2 | 3.0 | 2.3 | 2.0 | 1.3 | 0.8 | 2.2 | 2.3 | 20 |
| $P_{nn}$ | 0 | 0.57 | 0.02 | 0.17 | 0.11 | 0 | 0 | 0 | 0.02 | 0.11 | 1.0 |
| soft$P_{nn}$ | 0.07 | 0.57 | 0.33 | 0.46 | 0.40 | 0.11 | 0.03 | 0.03 | 0.40 | 0.32 | 2.7 |
| wE max | 0.25 | 3.2 | 1.5 | 4.2 | 2.2 | 0.43 | 0.04 | 0.02 | 2.2 | 1.5 | 15.5 |
| $\langle E \rangle$ | 0.29 | 0.67 | 1.6 | 0.75 | 1.2 | 0.055 | 0.09 | 0.1 | 0.57 | 0.79 | 5.8 |
| $\chi^2$ | 0 | 0.08 | 1 | 0 | 0.95 | 0 | 0 | 0 | 0 | 0.04 | 2.07 |
| wE avg | 0 | 0.05 | 1.55 | 0 | 1.17 | 0 | 0 | 0 | 0 | 0.03 | 2.8 |
| Expected max $\delta/\sigma$ | 3.5 | 2.3* | 2.8 | 3.3 | 2.7 | 3.4 | 3.4 | 2.8 | 2.7 | 2.8 | 31.6 |
| expected - actual max $\delta/\sigma$ | 1.7 | -0.07* | 0.6 | 0.27 | 0.4 | 1.4 | 2.1 | 2.0 | 0.4 | 0.6 | 11.5 |
| max E | 3.4 | 5.6 | 4.7 | 9.0 | 5.5 | 4.0 | 1.6 | 0.62 | 5.4 | 4.6 | 44.4 |
| expected max E | 12 | 5.3* | 7.7 | 11 | 7.5 | 12 | 12 | 7.8 | 7.4 | 7.7 | 102 |
| expected /actual E | 3.6 | 0.94* | 1.6 | 1.2 | 1.4 | 3.0 | 7.2 | 13 | 1.4 | 1.7 | 57.9 |
| expected<br>-actual E | 8.8 | -0.34* | 2.9 | 1.7 | 2.1 | 8.0 | 10 | 7.3 | 2.0 | 3.0 | 58.0 |

Note that only 31 non-bonds are listed because only 31 non-bonds had positive L-J energy values. Negative energies were neglected in the worst-outlier and average energy calculation for non-bonds, which were separate from the 12th E_LJ_ category. The lowest possible wE score is therefore −1.0. This would occur if all bonds, angles, torsions, and other chemical geometry in the structure are exactly equal to their ideal values, which is unlikely, but theoretically possible. There is no limit to how high wE can go, but it tends to plateau above 110 when there is at least one bad outlier > 3.3 σ in every category. Many clashes and bad non-bonds can push it even higher. In general, however, if all categories behave like Gaussian distributions and N_things_ is >1000 then wE ∼= 60 is expected. This is because the maximum expected deviates will all be clipped at max E∼10 by Equation 6, both weights (χ2 and P_nn_) wil be 0.5, the expected average squared Gaussian deviate (<E>) is 1.0 and there are 11 categories: 0.5*1.0*11+0.5*10.0*11 = 60.5. The ground truth model here is much better than this because it was made to have particularly good geometry. The Level 4 structure, however, has wE=54.6 because of the distortions required to fit a largely single-conformer model into this fundamentally 2-conformer density.

The exact code for calculating wE from a PDB file is released with the Phenix suite and also available as a shell script in the Challenge files.

## Appendix C Details of ground-truth model preparation

Beginning with the deposited structure and data for PDB id 1aho, refmac5 (Murshudov *et al*., 2011) was used to add missing atoms, including hydrogens. This did not include, by default, the terminal OH hydrogens of Ser, Thr and Tyr residues, which are commonly disordered. Each atom was then split into two conformers: A and B, starting at the C-terminus where a disulfide bridge with 2 conformations was clearly visible in the real electron density. Further splitting was performed manually using Coot (Emsley *et al*., 2010) until all 582 non-H atoms, including waters, had exactly two conformations. The structure was then optimized through a series of manipulations beginning with exhaustive rounds of single-atom swap-and re-refine (§3.4), selecting the best score as the starting point for the next round.

The scoring function used for this atom-swapping optimization was a rudimentary ancestor of the wE score described above. It consisted of the product of R_work_ with the molprobity score (MP), the average deviate/sigma (<Z<) in each of five categories: peptide omega angle, ramachandran, rotamer and C_β_ deviations, and non-bonded interactions. To this, in turn, was multiplied the worst sigma-scaled deviate (Z) in each of five categories: bonds, angles, dihedrals, chirality, and planarity.

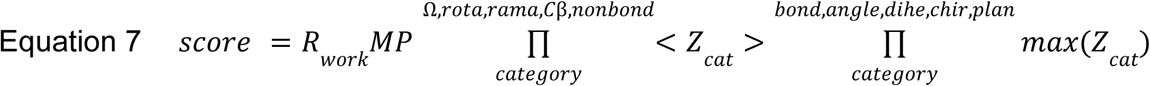

When the score converged for single atom swaps, pairs of atoms were then swapped until convergence. Convergence here is defined as arriving at a model where any and all further attempts to swap and re-refine always results in a worse score. Single and pair swaps were then iterated to convergence. The third step involved swapping groups of atoms: side chains, main chains, peptide bonds, and finally large stretches of sequence, again iterating to convergence. Finally, moves that had previously improved the energy score individually were tried in combination with each other. For the final, swap-optimized model, all further individual or combined conformer swaps with re-refinement were found to always give a worse score.

The swap-optimized model was then further refined using a targeted weight-modification approach using phenix.refine (Afonine *et al*., 2012). Non-default settings included wxc_scale=0.1, chir_volume_esd=0.1, c_beta_restraints=True and disabling features like occupancy refinement. The worst outlier bonds, angles, torsions, and other restrained entities were identified, the strength of the particular restraint was increased, and the refinement repeated from the start. Clashes were treated as bad non-bond contacts, and combatted by adding a new, artificial “crowbar” bond between clashing atoms that was specified to be longer than the gap it spans. The strength of this crowbar bond restraint was increased until the gap became open enough not to clash. This process was iterated with the worst outlier in each category eventually giving way to the second worst as a target for weight modification. Eventually, after 289 rounds, few significant geometry outliers existed, save for three torsion outliers: Cys12, Glu32 and Gln37. These were excluded from targeted weight modification because they are clearly visible as strained in the real electron density. A bad non-bond contact in the backbone of Glu32 that no doubt contributes to the Glu32 rotamer outlier was reduced by targeted restraints, but not eliminated.

The goal of this rather unorthodox refinement procedure was to generate a model that was roughly consistent with the real-world electron density of this protein, but free from chemical strain outliers larger than 3x standard deviation, which is often considered large enough to be statistically questionable. This was not intended to make the ground truth unrealistically “good”, but rather to be sure it was not arguably implausible. For example, if the ground truth contained a highly stretched bond for no particular reason, an algorithm that requires ensemble members to have good bond lengths might not be able to capture it.

Summary of restraint weight changes, and final worst-outliers in best.pdb:

| category | bond ang | Non-bond | pept $\Omega$ | side $\chi$ | Cβ dev | plane | bond len. | chir | rota | rama | sum |
| --- | --- | --- | --- | --- | --- | --- | --- | --- | --- | --- | --- |
| N values | 1420 | 30,656 | 126 | 628 | 114 | 1192 | 1046 | 134 | 108 | 124 | 4923 |
| modified | 34 | 9 | 29 | 18 | 15 | 3 | 0 | 0 | 0 | 0 | 108 |
| >2x tight | 2 | 8 | 17 | 15 | 3 | 1 | 0 | 0 | 0 | 0 | 46 |
| >4x tight | 0 | 0 | 0 | 2 | 0 | 1 | 0 | 0 | 0 | 0 | 3 |
| max $\delta/\sigma$ | 1.8 | (2.4) | 2.2 | 3.0 | 2.3 | 2.0 | 1.3 | 0.8 | 2.2 | 2.3 | 20 |
| location | C46 | E32 | A39 | E32 | V10 | Y14 | C16 | K30 | C12 | D8 | - |
| max E | 3.4 | 5.6 | 4.7 | 9.0 | 5.5 | 4.0 | 1.6 | 0.62 | 5.4 | 4.6 | 44.4 |
| <E> | 0.29 | -0.32 | 1.6 | 0.75 | 1.2 | 0.055 | 0.09 | 0.1 | 0.57 | 0.79 | 5.8 |

No changes were made to bond length, planarity, chirality, or Ramachandran restraints. Rotamers were indirectly impacted by side chain torsion restraints. Note that non-bond interactions here do not have a standard deviation *per se*, because the Lenard-Jones 6-12 energy is being used to score them. The square root of this energy is provided as the σ deviate in parentheses above for context.

The final wE geometry score (described above) of the ground-truth model was 11.8027, which is excellent. The phenix.fmodel bulk solvent parameters: k_sol=0.42 b_sol=45 solvent_radius=1.1 shrink_truncation_radius=0.9, were chosen for mutual compatibility with refmac5.

This test data set served the Challenge well, but it was not without flaws, which we list here for future reference. The original 1aho data set was missing 14% of the diffraction data, and these missing data were passed on into refme.mtz. These absent reflections led to significant skewness in the difference map, and a false peak at 5.1 σ. Although statistically unlikely for a Gaussian distribution, a skew Gaussian can support such apparent outliers. This hampered application of P_nn_-like scoring of density difference peaks. In the future, this will be addressed by a more general probability score, but also using data sets with 100% completeness. A bug in generating Gaussian errors from SIGF also produced some negative F values in refme.mtz. This did not appear to impact results, but will be avoided in the future by simulating diffraction patterns as a way to generate realistic errors, as in Figure 1.

